# Pioneering, chromatin remodeling, and epigenetic constraint in early T-cell gene regulation by PU.1

**DOI:** 10.1101/331710

**Authors:** Jonas Ungerbäck, Hiroyuki Hosokawa, Xun Wang, Tobias Strid, Brian A. Williams, Mikael Sigvardsson, Ellen V. Rothenberg

**Author notes:** Co-first authors, J. U. and H. H.

## Abstract

PU.1 is a dominant but transient regulator in early T-cell precursors and a potent transcriptional controller of developmentally important pro-T cell genes. Before T-lineage commitment, open chromatin is frequently occupied by PU.1, and many PU.1 sites lose accessibility when PU.1 is later downregulated. Pioneering activity of PU.1 was tested in in this developmentally dynamic context, by quantitating the relationships between PU.1 occupancy and site quality and accessibility as PU.1 levels naturally declined in pro-T cell development, and by using stage-specific gain and loss of function perturbations to relate binding to effects on target genes. PU.1 could bind closed genomic sites, but rapidly opened many of them, despite the absence of its frequent collaborators, C/EBP factors. The dynamic properties of PU.1 engagements implied that PU.1 binding affinity and concentration determine its occupancy choices, but with quantitative tradeoffs for occupancy between site sequence quality and stage-dependent site accessibility in chromatin. At non-promoter sites PU.1 binding criteria were more stringent than at promoters, and PU.1 was also much more effective as a transcriptional regulator at non-promoter sites where local chromatin accessibility depended on the presence of PU.1. Runx motifs and Runx1 binding were often linked to PU.1 at open sites, but highly expressed PU.1 could bind its sites without Runx1. Notably, closed chromatin presented a qualitative barrier to occupancy by the PU.1 DNA binding domain alone. Thus, effective pioneering at closed chromatin sites also depends on requirements beyond site recognition served by non-DNA binding domains of PU.1.

## Introduction

A central question in developmental molecular biology is how much a cell’s regulatory history affects the genomic sites where a given transcription factor can work, and under what conditions that factor can then remold the chromatin constraints under which future regulators will act. Different pre-existing chromatin accessibility landscapes can help to restrict binding so that the same factor may preferentially engage different regulatory elements (Treiber et al. 2010). However, questions remain about how much of a barrier chromatin state may pose, or whether there is a clear distinction between transcription factors that are highly chromatin-limited and other, “pioneer” factors, that might engage their sites impervious to chromatin state. PU.1 (*Spi1, Sfpi1*) is a transcription factor that works both to mediate developmental choices of blood progenitor cells and to serve the alternative developmental fates – associated with different epigenetic landscapes – that emerge from these choices. Its role in T-cell development is confined to the early stages leading up to T-cell lineage commitment, where it occupies tens of thousands of genomic sites in uncommitted cells but disappears in a programmed way during commitment. This dynamic transition offers a rare window into the quantitative contributions of affinity, chromatin context, factor concentration, and developmental history as predictors of PU.1 function.

The best studied roles for PU.1 involve its sustained actions as a lineage-determining transcription factor in myeloid, dendritic-cell, and B cell development, where it can open closed chromatin and recruit other transcription factors including C/EBPα, NF-κB, and IRF4 or 8 (Escalante et al. 2002; Carotta et al. 2010; Heinz et al. 2010; Natoli et al. 2011; Ostuni et al. 2013; McAndrew et al. 2016)[reviewed in (Gosselin and Glass 2014)]. In the macrophage context, PU.1 can act as a pioneer factor, capable of evicting nucleosomes (Barozzi et al. 2014), but collaboration with known partners also affects PU.1 site choices (Heinz et al. 2013). PU.1 is also crucial for multiple other hematopoietic lineages (Back et al. 2005; Iwasaki et al. 2005; Nutt et al. 2005) including the earliest stages of T-cell development (Champhekar et al. 2015).

Although most mature T cells express no PU.1, pro-T cells in the earliest stages maintain high PU.1 levels through multiple cell divisions before downregulating it during T-cell lineage commitment (Yui et al. 2010; Zhang et al. 2012). C/EBP or IRF family factors that PU.1 cooperates with in myeloid and B cells are expressed little if at all in early T-lineage cells. The lack of C/EBP factors should affect its genome-wide binding profiles, as shown by elegant analysis of natural genetic variation at collaborative binding sites (Heinz et al. 2013). Yet its potential for pioneering activity in other cell types, enhanced by its unusual protein stability (Kueh et al. 2013), could make it a strong contributor to “epigenetic memory” in early pro-T cells. Here we have investigated how it may work to set the chromatin context in which the T-cell program emerges.

PU.1 expression is well established in multilineage lymphoid precursors before they enter the thymus. Within the thymus, pro-T cells proliferate and traverse a sequence of phenotypically defined DN1, DN2a, DN2b, and DN3a stages before they express T-cell receptor proteins. They become committed to the T-cell fate only during the DN2a to DN2b transition (Fig. 1A), which occurs only after multiple cell divisions under the influence of thymic Notch pathway signaling [rev. in (Yui and Rothenberg 2014)]. This transition encompasses a global shift in 3-D chromatin associations (Hu et al. 2018) and in regulatory gene expression (Rothenberg et al. 2016), and this is when PU.1 expression is finally shut down. Until commitment, PU.1 binds to >30,000 sites in DN1 and DN2a pro-T cells, distinct from sites occupied in B and myeloid lineage cells (Zhang et al. 2012), and supports optimal proliferation and restrains specific alternative lineage genes, but slows progression toward commitment, thus enforcing correct timing of access to T-lineage genes (Champhekar et al. 2015). In previous studies, we identified impacts of PU.1 on specific, developmentally relevant genes by acute gain and loss of function perturbations (Anderson et al. 2002; Dionne et al. 2005; Franco et al. 2006; Del Real and Rothenberg 2013; Champhekar et al. 2015). The results showed that PU.1 represses as well as activates genes (Dionne et al. 2005; Franco et al. 2006; Del Real and Rothenberg 2013), antagonizing Notch signaling in early T-cell development and delaying activation of many T-cell genes until commitment. However, PU.1 binding sites globally appeared to be associated with active chromatin and actively expressed genes (Zhang et al. 2012). Thus, the actual relationships between PU.1 binding at different classes of sites and its regulatory function in normal early T cell development remained to be defined. Despite the power of PU.1 to open chromatin, most genes linked to PU.1 occupancy *in vivo* do not change expression as PU.1 levels decrease in T-cell development (Zhang et al. 2012): thus, most PU.1 binding events do not appear to control transcription. Moreover, because multiple “phase 1” transcription factors change expression in parallel with PU.1(Yui and Rothenberg 2014; Rothenberg et al. 2016), correlation alone cannot determine which developmental changes in gene expression are due to changes in PU.1 itself. Here, therefore, we have used gain and loss of function perturbations coupled with genome-wide analyses of the changes they induce in binding, chromatin state, and transcription of developmentally significant genes to show how PU.1 selects the sites that it will occupy in changing chromatin contexts, respecting or reshaping local chromatin states, and how its binding at different classes of sites controls the early T-cell program.

**Figure 1:**
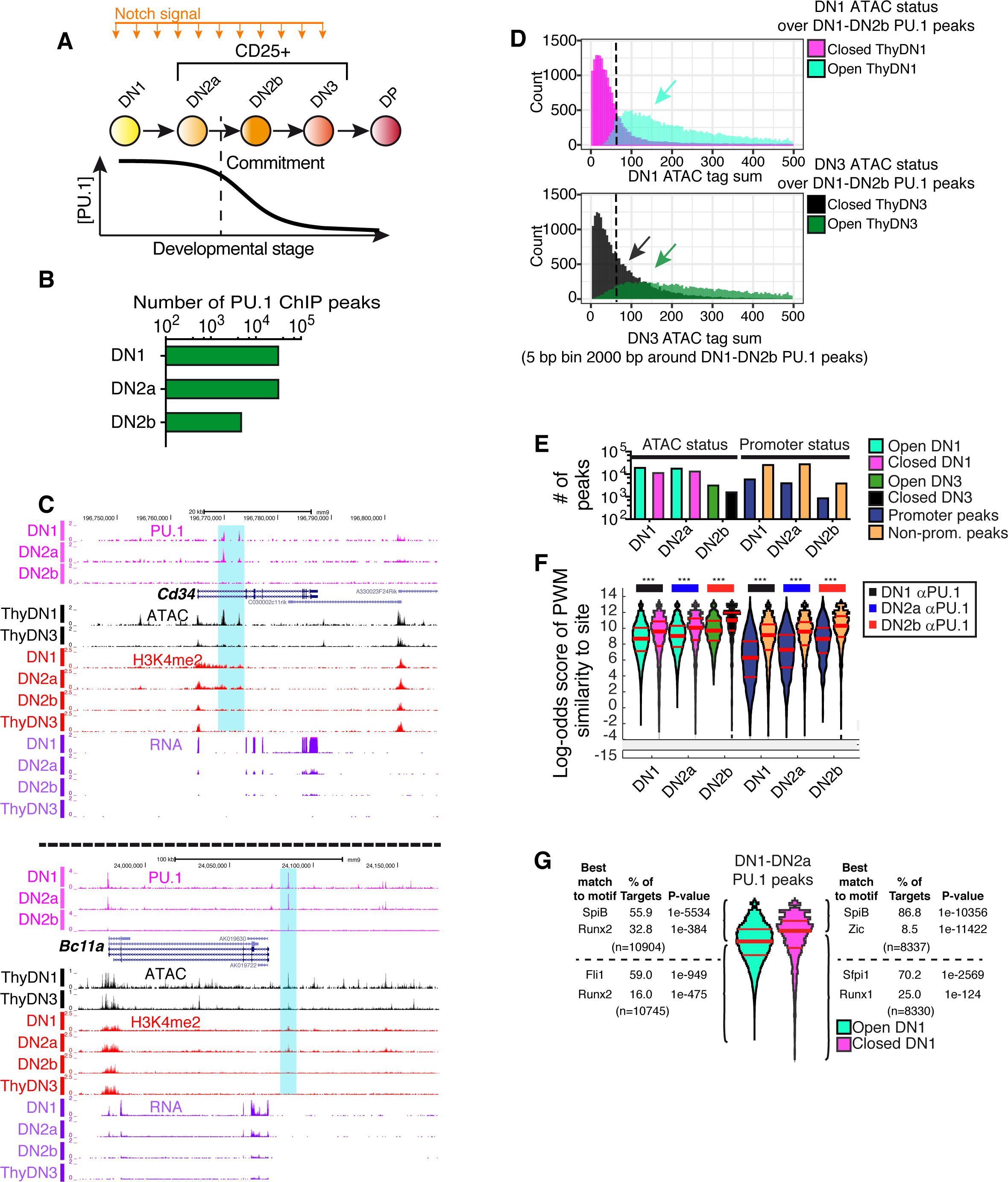
Endogenous PU.1 binding in pro-T cells is correlated with site accessibility and observes distinct affinity thresholds at open and closed sites and promoter and non-promoter sites. **A)** Outline of early T-cell development with schematic depicting PU.1 expression levels. **B)** Number of endogenous PU.1-occupancy peaks detected in DN1, DN2a, and DN2b pro-T cells. **C)** *Cd34* and *Bcl11a* UCSC browser tracks (genome.ucsc.edu) showing endogenous PU.1 ChIP, ATAC-seq, H3K4me2 ChIP, and RNA-seq in the DN1, DN2a, DN2b, and/or DN3 stages. Samples from in vitro differentiation from fetal liver precursors or from thymus (ThyDN1, ThyDN3). Data from (Zhang et al. 2012) except for ATAC (this study). **D)** Histogram of ATAC tag-counts at DN1 (top) or DN3 (bottom) stages, across all pro-T cell PU.1 binding sites. Sites in regions defined as ATAC ‘open’ or ‘closed’ (see Methods) are plotted separately to aid visualization. Y axis: number of sites at indicated ATAC signal level. **E)** Number of PU.1 peaks in indicated stages in ATAC-open and ATAC-closed as well as promoter and non-promoter regions. Same color key is used in (E) and (F). **F)** Distribution of DN1-DN2b motif log-odds scores at binding sites in open (cyan; open in DN1, green open in DN3) and closed regions (magenta closed in DN1, black; closed in DN3), and in promoters (dark blue) and non-promoter (orange) elements. Scores from a DN1-DN2b derived PU.1 PWM-matrix (Table S1). Kruskal-Wallis statistical test, *** *p* ≤ 0.0001. **G)** Motif analyses of PU.1 sites in DN1-DN2a cells, classified by PWM scoring and ATAC accessibility in ThyDN1.

## Results

### Dynamic PU.1 occupancy at open and closed chromatin sites in pre-commitment pro-T cells

PU.1 activity is a hallmark of the early, pre-commitment stages of T-cell development only (Fig. 1A). Accordingly, genome-wide, PU.1 occupancy decreases from about 30,000 sites of occupancy in DN1 and DN2a stages to ∼5000 sites in newly-committed DN2b cells (Fig. 1B), all distinct from sites in non-T lineage cells and the DN2b sites being a subset of the DN1-DN2a sites (Zhang et al. 2012). To evaluate how PU.1 might select these sites, we examined the chromatin status of PU.1 sites and the extent to which the binding specificity of the protein itself could determine its site preferences. This work used a sequence-agnostic criterion of regulatory site activity and took advantage of the natural downward titration of PU.1 during commitment (DN1 and DN2a to DN2b and DN3). Activity states of chromatin were mapped at high resolution as changes in DNA accessibility (“openness”) using Analysis of Transposase-Accessible Chromatin (ATAC)-seq (Buenrostro et al. 2013). Fig. 1C shows examples of PU.1 occupancy patterns at *Cd34* and *Bcl11a*, two target genes most highly expressed in pre-commitment DN1 cells, correlated with ATAC status of the relevant sites before (ThyDN1) and just after commitment (ThyDN3). Although little change in “permissive” H3K4me2 chromatin modifications occurs across these stages in general (Zhang et al. 2012), these loci are linked with specific sites where developmental loss of PU.1 binding during commitment is accompanied by losses of ATAC accessibility and H3K4me2 (Fig. 1C, highlighted regions). Considering all the sites genome-wide where PU.1 bound during any DN1-DN2b stage, ATAC criteria showed that more of these sites were likely to be “open” in DN1 stage (Fig. 1D, cyan) than in DN3 stage, when PU.1 was almost gone (Fig. 1D, dark green). This suggests that PU.1 either causes these sites to open or is preferentially recruited to them because of their accessibility.

Changes in ATAC status of PU.1 sites between DN1 and DN3 stage were strongly correlated with changes in expression of the closest linked gene. As shown by an empirical cumulative distribution frequency (ECDF) plot (Fig. S1A), genes linked to PU.1 sites that lost accessibility during commitment (“closing sites”) were more likely to be downregulated (DN3/DN2a ratio <1, padj<0.1, blue arrow) than genes without PU.1 sites. Accordingly, genes linked to sites that increased accessibility (“opening sites”) were more likely to be upregulated (DN3/DN2a ratio >1, padj<0.1, red arrow). Taken together with the enrichment of open regions among all PU.1-occupied sites, this suggests that PU.1 could positively regulate hundreds of genes specific to precommitment cells. However, PU.1 binding across the genome was not confined to open sites (Fig. 1D). In fact, in DN1 and DN2a cells, PU.1 occupied similar numbers of closed and open sites (Fig. 1E). Therefore, PU.1 does not simply engage sites epigenetically primed by other factors, nor does its binding always cause chromatin opening.

### Distinct PU.1 affinity thresholds in open and closed chromatin and at promoter and non-promoter sites

In macrophages, PU.1 has been established as a pioneer factor capable of displacing nucleosomes from its binding sites (Barozzi et al. 2014). In pro-T cells, where PU.1 expression is inherited from pre-thymic precursors but its best-characterized partner factors are absent, it was not clear whether PU.1 led or followed other factors to select its sites. The definition of pioneering (Zaret and Carroll 2011) implies that the DNA binding specificity of the factor itself guides its binding to sites in initially closed chromatin. We tested whether PU.1 intrinsic specificity guides its binding to “open” and “closed” sites by scoring PU.1 occupancy sites genome-wide for log-odds match to an optimal position weight matrix (PWM), calculated *de novo* (Table S1, Fig. S1B) from its pro-T cell binding sites (Zhang et al. 2012). The scoring pattern of sites across the genome (Fig. 1E,F) agreed well with the pattern scored using a more permissive PWM, calculated from PU.1 sites occupied in macrophages, which was reported to reflect PU.1 binding preferences on naked DNA (Pham et al. 2013)(Fig. S1B,C). The canonical PU.1 motif was most enriched at all classes of sites (Fig. S1D). However, at each stage, PU.1-occupied sites in open regions had a broader range of quality scores, with lower median and quartile quality scores, than the PU.1-bound sites in closed regions, implying that higher affinity was needed to establish binding at closed sites. The “closed” PU.1 binding sites accordingly showed a greater enrichment of a PU.1 subfamily (Sfpi1, SpiB) ETS motif than the “open” sites (Figs. 1G, S1D). On the other hand, open sites generally as well as the lower-quality half of the closed sites were distinguished from higher-quality closed sites by their high co-enrichment of Runx family motifs (Figs. 1G, S1D).

The changes in PU.1 occupancy across the genome as PU.1 levels naturally decrease in development implied that PU.1 binding site selection is governed by its own mass action. As PU.1 tag counts at individual sites and the number of scorable binding regions genome-wide both dropped during commitment (Fig. 1B), the sites still occupied tended to be those with higher motif quality (Figs. 1F; S1C,E). However, distinct binding criteria applied in open and closed chromatin regions. Both open and closed regions lost PU.1 occupancy (Fig. 1E, left), but the site quality difference between them persisted where PU.1 remained bound (Fig. 1F, left), consistent with a continuing affinity “penalty” in the closed regions. In the same cell nuclei, declining PU.1 levels also yielded different effects on occupancy affinity thresholds at promoter and non-promoter (distal) sites. Promoter sites were occupied even when they had poor quality target motifs (Fig. 1F, right), and when PU.1 fell to lower levels, poorer quality promoter sites remained occupied than non-promoter sites. This global difference between criteria for binding at promoter and non-promoter sites was evident at each developmental stage, and even when open and closed promoter and non-promoter sites were compared separately (Fig. S1E). Thus, PU.1 is recruited differently to promoters and enhancer elements.

### Non-promoter PU.1 sites control chromatin configuration and function

PU.1 binding does not always open chromatin, but does the accessibility of its open binding sites depend on continuing PU.1 occupancy? To test this, we first correlated genome-wide PU.1 binding dynamics with changes in local chromatin status. The ∼40K sites of PU.1 occupancy in the genome were clustered based on PU.1 binding, ATAC accessibility, and H3K4me2 modification status from DN1 to DN2b or DN3 stages as shown in Fig. 2A,B. The majority of sites fell into either of two major groups (Fig. 2A, Group 1 and Group 2; K-means). About 60% of all sites of PU.1 occupancy in DN1 and DN2a cells lost accessibility, and sometimes even the “permissive” histone mark H3K4me2, as PU.1 expression declined in DN2b/DN3 stages (Fig. 2A,B; site Group 2, “closing”). The other large group of sites were mostly open sites that stayed highly accessible and independent of developmental stage or level of PU.1 binding (Fig. 2A,B, Group 1, “static”). In contrast, no distinct subgroup of sites that “opened” as PU.1 levels declined was salient enough to emerge using a variety of clustering algorithms. Thus, the dominant division of PU.1 occupancies was into sites with constitutive accessibility and sites whose accessibility decreased in parallel with PU.1 expression itself.

**Figure 2:**
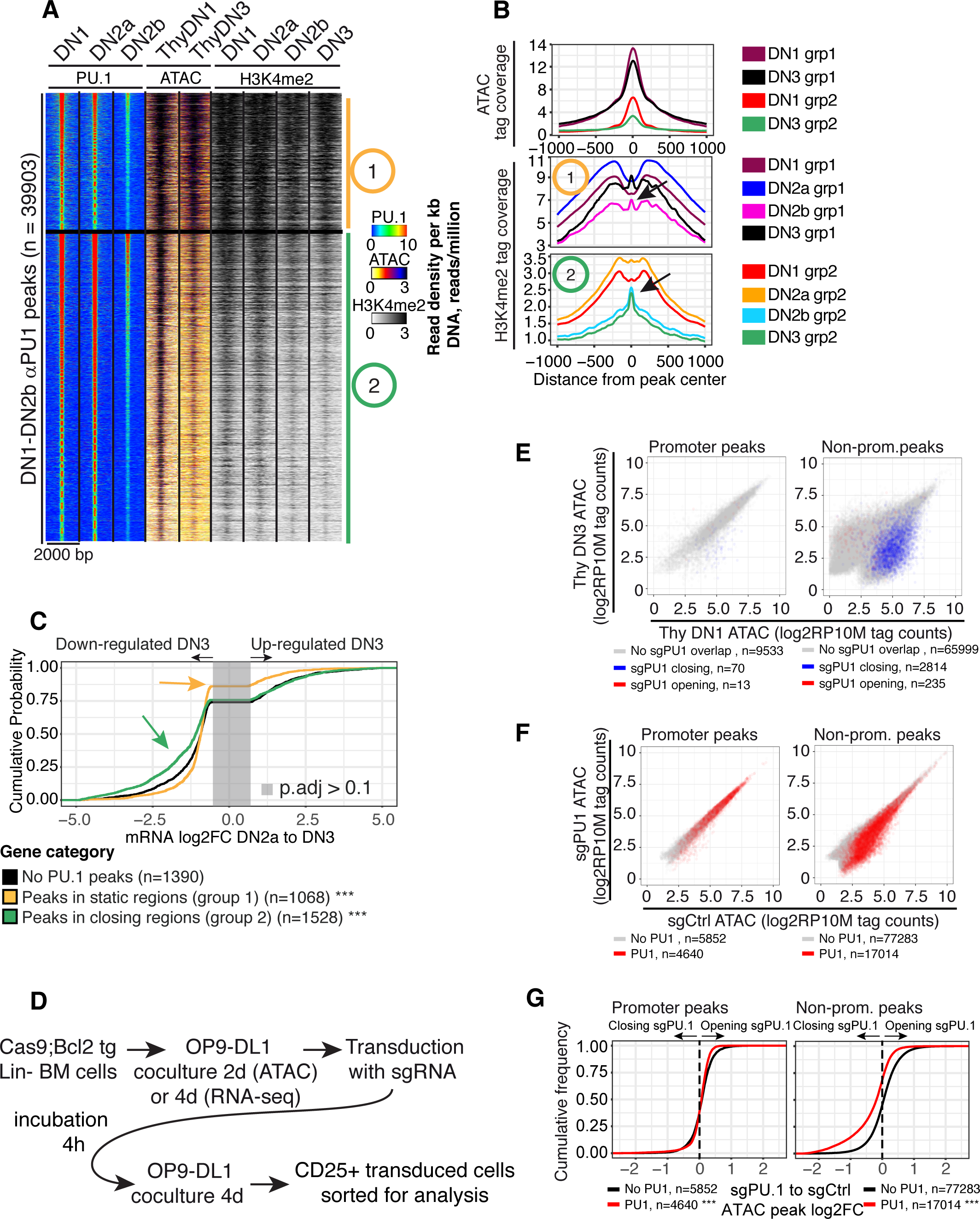
Loss of PU.1 is associated with decreased chromatin accessibility and loss of transcriptional activity. **A)** Heatmap of PU.1 and ATAC tag count distribution in early DN-pro-T development. Two kb-wide regions were K-means clustered (K=2) based on PU.1 binding, ATAC, and H3K4me2 patterns. Constitutively open sites (Group 1) and sites losing accessibility from DN1 to DN3 (Group 2) are distinguished. **B)** ATAC and H3K4me2 distributions at the indicated stages at sites in Groups 1 and 2 in **A**. Arrows in H3K4me2 plot: nucleosome re-constitution after PU.1 loss. **C)** Natural changes in gene expression across commitment (DN2a to DN3) linked to PU.1 bound sites from Fig 1F Groups 1 (static) and 2 (closing). *** Kolmogorov-Smirnov p-value ≤ 0.0001. **D**) Schematic of protocol for Cas9-mediated disruption of PU.1 using transduction with sgPU.1 or control guide RNAs. **E)** Effects of disruption of PU.1 in DN2 cells on accessibility of promoter and non-promoter sites. X axis: ATAC accessibility of individual genomic sites in DN1 stage; y axis, accessibility in DN3 stage. Blue: sites that lose accessibility upon knock-out of PU.1 in DN2 cells; red: (rare) sites that gain accessibility. **F)** Enrichment of PU.1-binding sites among genomic sites that lose ATAC accessibility upon PU.1 deletion (see panel **D**). ATAC-accessibility of genomic sites after PU.1 deletion (y axis) is plotted against their accessibility in controls (x axis). Sites overlapping with PU.1 binding in DN1, DN2a, and/or DN2b cells are displayed in red. **G)** Association of PU.1 binding with loss of accessibility upon PU.1 knock-out described in E. *** K-S p-value ≤ 0.0001.

PU.1 binding at non-promoter sites was correlated with specifically pre-commitment gene expression. Genes linked only to Group 1 binding sites changed little in expression between DN2a (high PU.1) and DN3 (low PU.1) stages (Fig. 2C), even less (yellow arrow, p<0.0001) than genes without any PU.1 binding peaks at all (Fig. 2C, black curve). In contrast, genes linked to Group 2 PU.1 sites (Fig. 2C green) changed expression much more than genes without peaks, most of them losing expression in this developmental transition (green arrow, p<0.0001, skew to left). This was not due to differential loss or retention of PU.1 between Group 1 and 2 sites (Fig. S1F). However, the static sites (Group 1) were highly enriched for annotated promoters (39.8% in promoter regions)(χ^2^ test p <0.0001), whereas the dynamically closing sites (Group 2) were overwhelmingly in non-promoter elements (4.6% peaks in promoter regions). These results suggest that PU.1 binding in pro-T cells is more stringent, more important for chromatin site accessibility, and better correlated with transcription when it binds at non-promoter regions than at promoters.

To test whether PU.1 itself was responsible for the dynamics of ATAC accessibility at these non-promoter sites, we exploited the Cas9 transgenic mouse strain (Platt et al. 2014) to provide T-cell precursors in which PU.1 could be acutely deleted at a specific developmental stage (Fig 2D). Hematopoietic precursors from these mice were isolated and induced to begin T-lineage differentiation *in vitro* by coculture with OP9-DL1 stromal cells. When PU.1 was at its height during DN1 to DN2a stages, specific guide RNAs were introduced by retroviral transduction (sgPU.1 or sgControl), and effects on chromatin accessibility were assayed by ATAC-seq after four more days of culture. This protocol deletes targeted exons biallelically (Fig. S1G) and removes endogenous PU.1 expression (Hosokawa et al. 2018). While promoter sites were unaffected, far more non-promoter sites specifically lost accessibility than gained accessibility when PU.1 was deleted, and these sensitive sites were normally open in DN1 but not in DN3 stage (Fig. 2E). These sites with PU.1-dependent accessibility were indeed direct targets, for they were also significantly enriched among distal sites that normally bound PU.1 in control DN1 cells (Fig. 2F,G; p ≤ 0.0001). The whole accessibility distribution of PU.1 binding sites in non-promoter regions shifted toward closure when *Spi1* was disrupted (Fig. 2G, right). In contrast, PU.1 deletion had little if any effect on promoter sites, with or without PU.1 binding (Fig. 2E-G). Thus, sustained PU.1 activity is needed to maintain ATAC accessibility at many non-promoter PU.1 binding sites and PU.1 is a primary determinant for open chromatin at these elements.

### Pioneering by PU.1 in DN3-like cells: site choice and chromatin opening in a closed epigenetic landscape

PU.1 not only maintained the open status of its normal binding sites but could also cause sites to open in closed chromatin. We used the Scid.adh.2C2 model cell line (Dionne et al. 2005) to measure this in a defined, low-background pro-T-cell context. These are immortal, clonal DN3-like cells that do not express endogenous PU.1, but respond sensitively to introduction of exogenous PU.1 with responses similar to those of primary pro-T cells (Dionne et al. 2005; Del Real and Rothenberg 2013)(Fig 3A). To place PU.1 activity under tight temporal control, Scid.adh.2C2 cells were transduced stably with full-length PU.1 linked to the ligand-binding domain of a tamoxifen-dependent estrogen receptor (PU.1ert2). Most of the PU.1 expressed in these cells is tethered in the cytoplasm, where large reservoirs can accumulate without affecting phenotype, but the PU.1 can translocate quickly and synchronously to the nucleus in response to 4-hydroxytamoxifen (4-OHT), causing cell-surface CD25 downregulation beginning by 24h (Fig. S2A), and CD11b upregulation later (Dionne et al. 2005; Del Real and Rothenberg 2013). From 0 to 24h of stimulation with 0.1 μM 4-OHT, we compared the kinetics of PU.1 binding to the genome with changes in H3K27Ac, ATAC accessibility, and RNA expression linked to the new binding sites (Fig. 3B, C; Fig. S2B), and found rapid occupancy at two large classes of sites (Group 3 and Group 4). Baselines were defined by control cells transduced with an Ert2-only control vector (“EV”, “EVert2”)(Fig. 3B,C)(gene expression list Table S2). The time course of binding had some background signal: in Scid.adh.2C2 cells stably transduced with PU.1ert2, low levels of PU.1ert2 reached the nucleus even without 4-OHT (PU.1ert2 at 0h of 4-OHT; Fig. S2C). Sites occupied in the presence of these low levels of nuclear PU.1 were restricted to already-active chromatin sites as judged by ATAC accessibility and H3K27Ac in non-transduced Scid.adh.2C2 or EV-transduced cells (Fig. 3B, Group 4; Fig. S2B,C). However, 4-OHT treatment induced nearly 5x more PU.1 binding sites within 2h (Fig. S2B,C), and most of this new binding was to Group 3 sites (Fig. 3B) that had no PU.1ert2 binding at 0h and were closed in unperturbed Scid.adh.2C2 cells (Fig. 3C, Fig. S2B,C). Examples are the highlighted sites in the *Icam1* gene, which were also bound by endogenous PU.1 in DN1-DN2a cells (Fig. 3D). Like Group 1 sites in primary cells (Fig. 2A), 51% of Group 4 sites were at promoters, contrasting with only 3.1% of Group 3 sites (χ^2^ *p*-value < 0.0001), and the PWM match quality for Group 3 sites bound by PU.1ert2 was much higher than for Group 4 sites (Fig. S2D).

**Figure 3:**
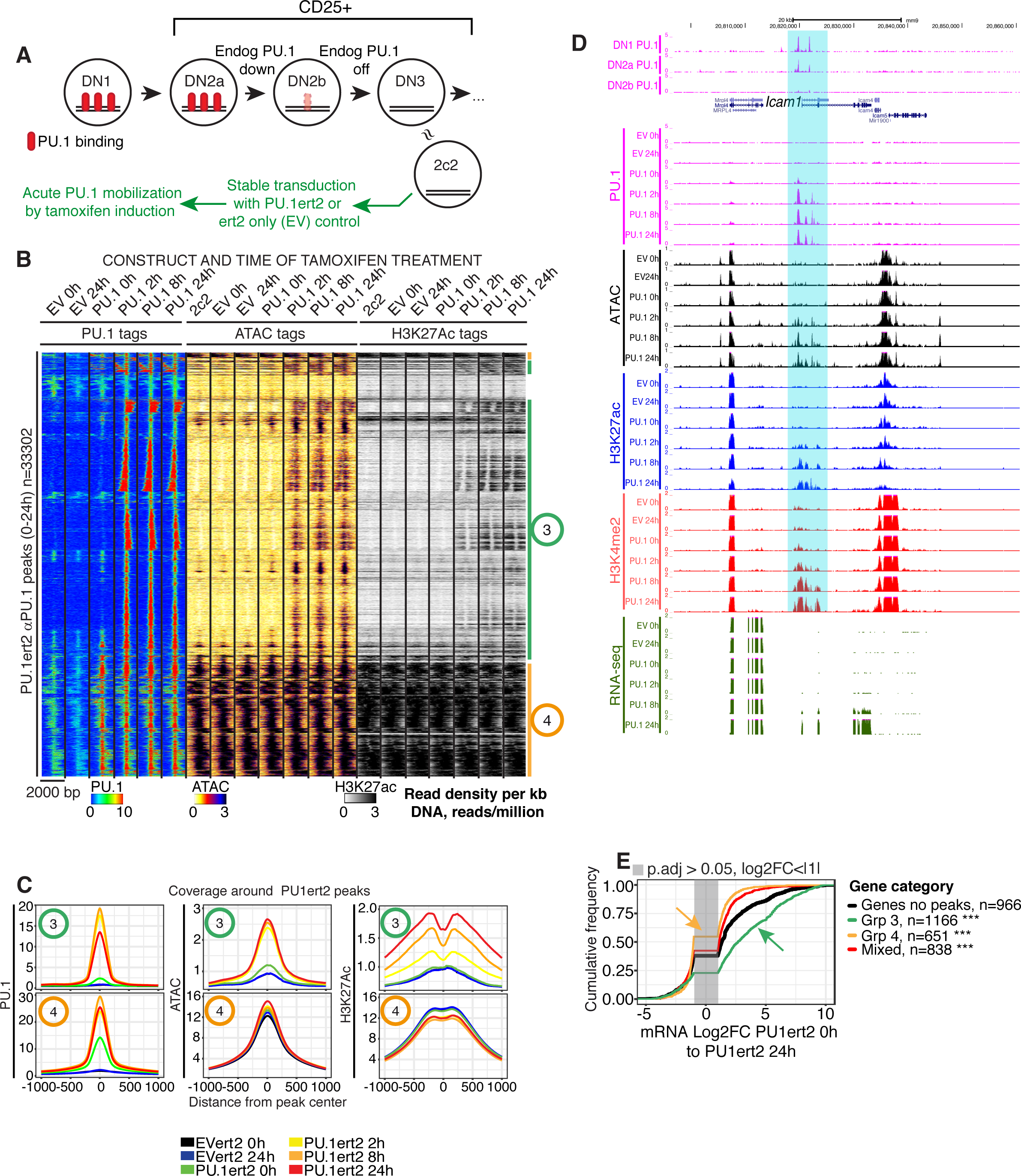
Short term induction of PU.1 activity leads to chromatin opening, altered histone modifications and cis-regulatory element-dependent gene activation. **A)** Schematic of experiment showing approximate relationship of Scid.adh.2C2 cells to normal program of T-cell development. **B)** Comparison of time courses of PU.1 binding and induced changes in chromatin accessibility and histone H3K27 acetylation after 0-24h of mobilization by 4-hydroxytamoxifen (4-OHT), in Scid.adh2C2 stably transfected with PU.1ert2 or EVert2. Sites in heat map were hierarchically clustered on Pearson correlation with average linkage. Shown are manually derived groups of sites that open and gain H3K27ac upon PU.1ert2 mobilization (Group 3) or that are already open and associated with H3K27ac at 0h (Group 4). **C)** Quantitative distribution plots of PU.1, ATAC and H3K27ac signals within 1 kb of PU.1 bound sites from Groups 3 and 4. **D)** Browser plots of PU.1 binding to the positively regulated *Icam1* gene in DN1-DN2b cells (top) and samples from the time course shown in Fig. 3B. Highlight indicates chromatin regions opened by PU.1 binding prior to RNA expression. **E)** Association of PU.1ert2 binding site Groups 3 & 4 with changes in linked gene expression between 0h and 24h of 4-OHT induction. *** K-S p-value ≤ 0.0001.

In the naïve context of this cell line, PU.1 exerted its main positive functional impact at closed, non-promoter Group 3 sites, rapidly making them ATAC accessible. This response was maximal already within 2h (Figs. 3B,C; Fig. S2B). Somewhat more slowly, ∼50% of the Group 3 sites also acquired the active histone mark H3K27Ac de novo, beginning at 2 h but with most signal observed by 8-24h of PU.1 binding (Fig. 3B,C; Fig. S2B). Genome-wide, genes responding to PU.1ert2 binding at linked Group 3 sites also usually increased expression over the 24h period (Fig. 3E, green). In contrast, those linked only to binding at Group 4 sites showed even less increase than genes without closely linked PU.1 sites at all (Fig. 3E, yellow).

Many functionally important PU.1 sites were bound immediately: the sites occupied at 2h were as likely to predict expression of linked genes at 24h as sites that were occupied only at 8h or 24h (Fig. S3A). However, the kinetics of transcriptional response implied a stepwise process. The pattern of expression of DEGs closely linked to new PU.1ert2 sites began to change from 2h onward (Fig. S3B), but only about 1/3 of the genes that were eventually activated showed upregulation by 8h, despite maximal ATAC accessibility already, and most were only activated by 24h (all p<10^−9^; Fig. S3B). Thus, PU.1 binding rapidly initiated local ATAC accessibility in previously closed chromatin sites enabling further steps resulting in local target gene activation. This is consistent with a pioneering mechanism for PU.1.

Despite the chromatin opening at most sites where PU.1 established binding, and despite the transcriptional activation that predominated at linked genes, PU.1 introduction also repressed genes in these cells as previously reported (Dionne et al. 2005; Del Real and Rothenberg 2013). Fig. S3B shows that the repression could start early, at least as early as most positive target genes were upregulated (also see Table S2). *Il7r* was substantially repressed even as PU.1 bound to an open site at its promoter (Fig. S3C). To evaluate activation vs. repression among PU.1’s natural roles, we applied perturbation tests to the primary-cell context.

### Gain and loss of PU.1 expression defines functional target genes in primary pro-T cells

To characterize the mechanisms of different physiological PU.1-mediated effects in pro-T cells, it was important to begin with the best comprehensive list of pro-T cell genes that normally respond to PU.1. We took advantage both of the good surface markers that identify different stages of T-cell development prospectively, and also of the efficient cell culture system that drives primary hematopoietic cell precursors through early T-cell differentiation. In this system, acute gain and loss of function perturbations of T-cell development can be carried out and analyzed at specific stages at will (Fig. 4A). In this system, to define the genes affected by acute gains and losses of PU.1 activity, we introduced exogenous PU.1 into differentiating, post-commitment pro-T cells and used CRISPR to delete PU.1 in pre-commitment pro-T cells, determining the global set of PU.1-dependent RNA expression changes.

**Figure 4:**
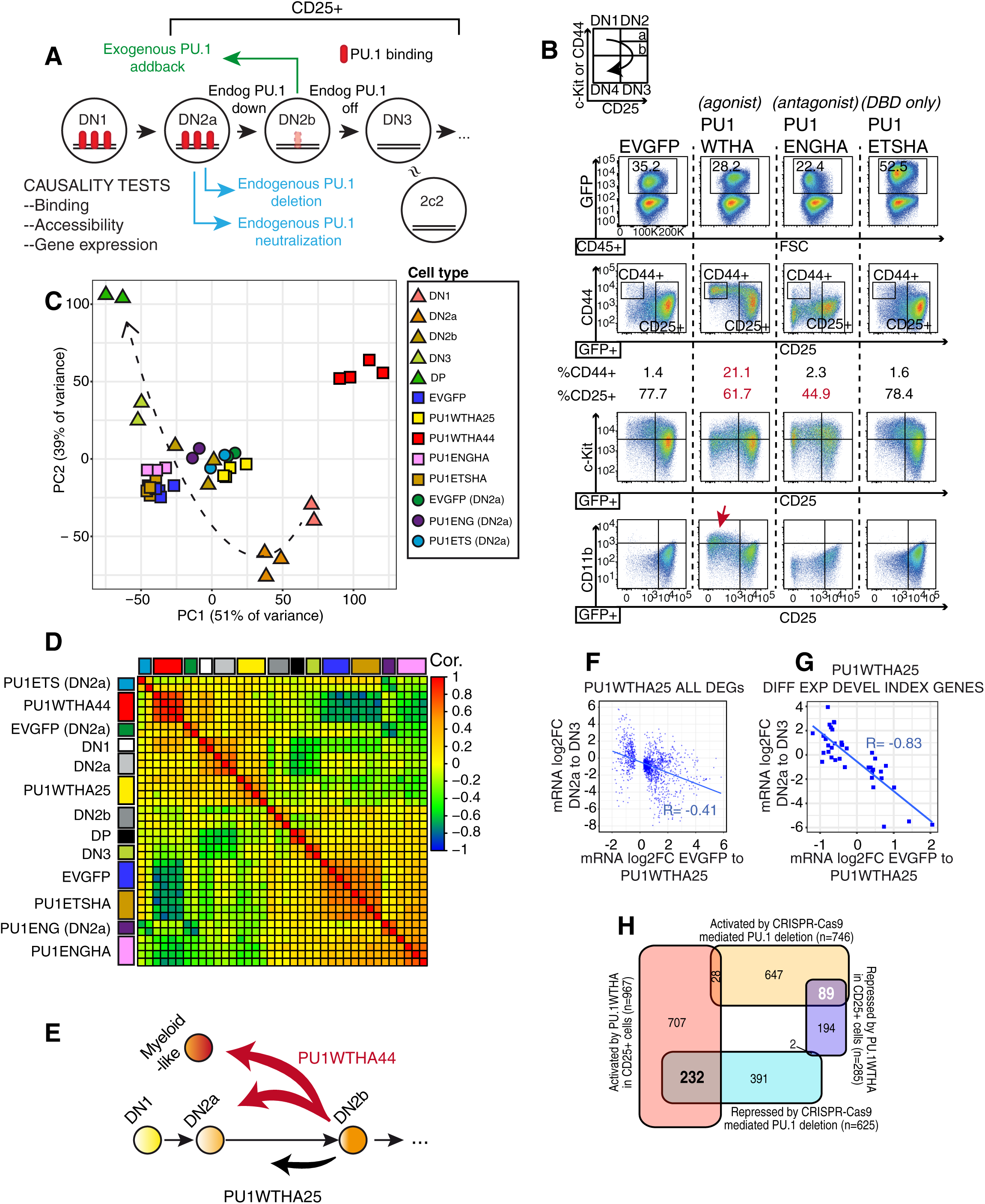
PU.1 introduction in CD25+ cells partly restores the earliest T-cell program. **A)** Schematic of acute gain-of-function and loss-of-function tests for causal role for PU.1 in gene expression changes during commitment. Colored arrows depict effects relative to normal development. **B)** Effects on developmental marker expression by reintroduction of PU.1, PU.1 antagonists, or empty vector (EV) into fetal liver-derived CD25+ (DN2b/DN3) pro-T cells. FACS analyses are shown 40h after transduction. Data are representative of four independent experiments. Insert: Normal developmental trajectory. **C)** Principal component analysis of expressed genes (RPKM ≥ 1) in four independent samples of CD25+ DN2b/DN3 cells transduced with EVGFP, PU1WTHA, PU1ENGHA, or PU1ETSHA, compared with DN2a PU.1 antagonist samples (GSE65344)(Champhekar et al. 2015), and normal reference cells (Zhang et al. 2012). Dashed arrow: approximate trajectory of normal development. **D)** Correlation analysis of gene expression as described in 4C. **E)** Schematic, effects of PU1WTHA on developmental gene expression in cells remaining CD25^+^ (PU1WTHA25) and cells becoming CD25-low CD44^+^ (PU1WTHA44). **F)** Transcriptional changes upon transduction of CD25+ cells with PU1WTHA as opposed to EV (blue) compared with normal transcriptional changes from DN2a to DN3 stages. Pearson’s r shown. **G)** Same as (**F**), for regulatory genes from developmental index list (Fig. S4D, Table S4). **H)** Venn diagram of genes differentially expressed after PU.1 gain of function (PU1WTHA25 vs. EV) and after Cas9-mediated deletion in pro-T cells (protocol in Fig. 2D). High-confidence targets with reciprocal responses to PU.1 loss and gain of function are indicated with number in bold face.

To enable observed gene expression changes to be related to occupancies, we used an epitope-tagged version of PU.1 (PU1WTHA) for gain of function. This construct, or a control empty vector (EV), was retrovirally transduced into pro-T cells generated from fetal liver progenitors, differentiated on OP9-DL1 stroma to DN2b-DN3 stage, i.e., when endogenous PU.1 was newly downregulated (Fig. 4A). A truncated PU.1 consisting only of the ETS DNA binding domain (PU1ETSHA), capable of competitively blocking some myeloid-lineage effects of endogenous PU.1 (Kueh et al. 2013; Champhekar et al. 2015), was also tested. Finally, to test whether PU.1 was really primarily an activator, we also compared effects of PU1WTHA with those of an HA-tagged obligate repressor form of PU.1 in which the ETS domain was fused to an Engrailed repression domain (PU1ENGHA)(Champhekar et al. 2015)(“endogenous PU.1 neutralization”, Fig. 4A). All were expressed at the protein level in the transduced cells by 40 h (Fig. S4A). Exogenous PU.1 and PU.1 antagonist constructs showed their typical effects (Del Real and Rothenberg 2013; Champhekar et al. 2015) by visibly altering the normal T-cell developmental progression. Early T-cell precursors progress from DN1 (Kit^+^ CD44^+^ CD25^−^, a.k.a. “ETP”) to DN2a (Kit^++^ CD44^+^ CD25^+^), through commitment to DN2b (Kit^+/int^ CD44^+^ CD25^+^) and then DN3 (Kit^−^ CD44^−^ CD25^+^)(Fig. 4B inset), and the PU.1 levels change most from DN2a to DN3 (all CD25+ stages; Fig. 4A,B). Whereas DN2b/DN3 cells transduced with empty vector (EV) progressed toward DN4 stage (CD25^−^ CD44^−^), PU1WTHA appeared to make them reverse direction partially (Fig. 4B, PU1WTHA vs. EVGFP), shifting toward DN1 or DN2a-like (higher CD44, many with loss of CD25). In contrast, cells transduced with PU1ETSHA resembled EVGFP controls, while the antagonist PU1ENGHA had opposite effects from PU1WTHA, decreasing CD44 expression (Champhekar et al. 2015).

To analyze direct effects of PU.1 itself, we sorted for the transduced cells that remained within the T-cell program. This was important because PU.1 not only works within the early T-cell program but also can promote lineage reprogramming, triggering other indirect effects (Anderson et al. 2002; Dionne et al. 2005; Lefebvre et al. 2005; Franco et al. 2006; Laiosa et al. 2006). Cells with moderate overexpression of PU.1 (∼3x above normal levels) were still CD25^+^ (“PU1WTHA25”), but cells with the highest PU.1 became CD44^+^ CD25^−^ lineage-diverted cells (“PU1WTHA44”), with low Kit expression and upregulated CD11b (*Itgam*)(Del Real and Rothenberg 2013). Comparison of the gene expression changes in PU1WTHA25 and PU1WTHA44 cells separately at 40 h post-transduction showed that the impacts of PU.1 were most likely to be direct in the PU1WTHA25 cells. RNA-seq (Table S3) and Principal Component Analysis of the samples (Fig. 4C) showed that the CD25^+^ cells in which we had perturbed PU.1 activity (yellow, brown, orange squares) remained generally close to controls (navy squares), and similar to DN2b cells differentiating from non-manipulated fetal liver precursors (Fig. 4C, triangles). These samples were also globally similar to samples transduced with the PU.1 antagonists for a shorter time in a previous study (circles)(Champhekar et al. 2015). In contrast, at this timepoint, PU1WTHA44 cells (red squares) had shifted far away from the pro-T cell trajectory. Fig. S4B,C shows there were numerous changes in other regulators in the PU1WTHA44 cells specifically that could contribute to this deviation. PU1WTHA44 cells repressed the essential Notch signaling pathway (*Notch1, Notch3, Hes1, Dtx1*, and *Nrarp*) as well as T-cell regulatory genes like *Bcl11b, Gfi1, Myb, Tcf7* and *Lef1* that were unaffected in PU1WTHA25 (Fig. S4B,C; Table S3). Uniquely also, PU1WTHA44 samples markedly upregulated myeloid specification genes including *Cebpa* and *Cebpd* and multiple IRF factor coding genes (Fig. S4B,C; Table S3), which were minimally induced in the PU1WT25 cells. The more selective gene expression changes in the PU1WTHA25 cells, therefore, were more likely to indicate effects of PU.1 dosage *per se* within the T-cell pathway.

Introduction of PU1WTHA into DN2b cells to convert them into PU1WTHA25 cells was indeed sufficient to shift their gene expression pattern “backward” toward a more pre-commitment, DN2a-like state (Fig. 4D, E), tending to reverse the effects of the normal DN2a-DN3 transition (Fig. 4F). This effect was more pronounced when evaluated on a developmental index list of 171 transcriptional regulatory genes that have robust, well-defined expression patterns in early T-cell development (Champhekar et al. 2015)(Table S4). Of these genes, 39 altered expression in PU1WTHA25 cells (qval <0.05; Fig. S4D), primarily by reversing their normal trends during DN2a to DN3 progression (Fig. 4G; Pearson’s r ≅ −0.83). Whereas effects on these genes too were often stronger in PU1WTHA44 cells, the indirect effects in those cells (Fig. 4E) made it preferable to focus on PU1WTHA25 cells for causal analysis. Thus, gain of function identified a set of PU.1-sensitive target genes intrinsic to the T-cell program which are also likely to be physiologically relevant targets of endogenous PU.1 in normal development.

### Cas9 deletion confirms shared targets of endogenous and exogenous PU.1

Genes responding to exogenous PU.1WTHA were indeed dependent on endogenous PU.1 normally for their positive or negative developmental control (Fig. 4A; Table S5), as shown using acute deletion in the Cas9 transgenic mouse system as described above (Fig. 2D)(Hosokawa et al. 2018). Due to the stability of PU.1 protein (Kueh et al. 2013), acute deletion effects are manifest with slower kinetics (days) than effects of introducing an exogenous wildtype or mutant PU.1 (<20 hr)(Champhekar et al. 2015). Therefore, we used adult bone marrow progenitors differentiating in OP9-DL1 culture, which progress more slowly than thymocytes or fetal liver-derived precursors between the pre-thymic stages where PU.1 is needed for viability and the post-commitment stages where PU.1 is no longer active (Huang et al. 2005).

Genes differentially expressed in these PU.1-disrupted cells showed highly biased patterns of overlap with those differentially expressed in PU1WTHA25 cells (Fig. 4H). Of the genes affected by both perturbations, >90% showed reciprocal responses (padj<0.05) to gain of PU.1 function in fetal-liver derived pro-T cells and to loss of PU.1 function in bone marrow-derived pro-T cells. Over 230 PU.1-dependent genes were concordantly induced by exogenous PU.1WTHA in FL-derived DN2b cells and inhibited by endogenous PU.1 deletion in the BM-derived DN2a/b cells, whereas almost 90 were concordantly repressed by exogenous PU.1 and upregulated by PU.1 deletion (Fig. 4H; Table S6A; extended list shown in Table S6B). These constituted a high-confidence list of the genes controlled by PU.1 in early T cells.

### Binding sites for exogenous PU.1 are preferentially linked to transcriptional activation in pro-T cells

Newly introduced PU.1WTHA established occupancy, determined by ChIP-seq in the CD25^+^ cells, at sites linked to >80% of the genes responding (padj<0.1) to PU.1 reintroduction in PU1WTHA25 cells. Of the highest-confidence targets (Table S6A, padj<0.05 in reciprocal perturbations), PU.1-occupied sites were linked to 90% of both PU.1-dependent genes and PU.1-constrained genes. The exogenous PU.1 in these DN2b-derived CD25^+^ cells efficiently reoccupied many of the same sites that had previously been occupied by endogenous PU.1 in DN1 and DN2a cells. Examples are shown at *Hhex* (Fig. 5A), which was strongly reactivated in post-commitment cells by exogenous PU.1. As most responding target genes were each linked to multiple sites of occupancy, however, we used a progression of queries to identify the most distinctive features of the particular sites that were likely to be important for function.

**Figure 5:**
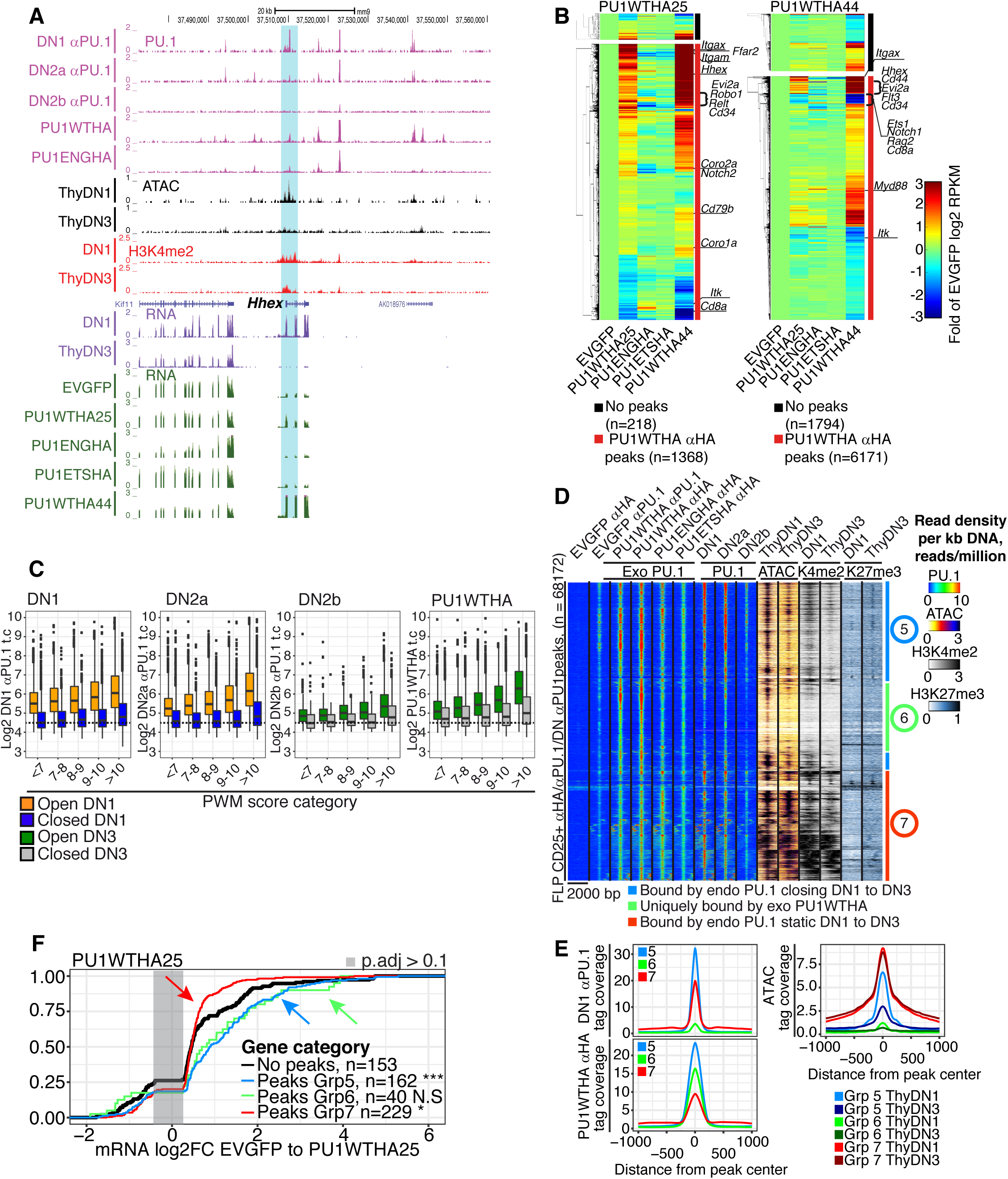
Structure-function relationships of sites of occupancy by exogenous PU.1 in primary pro-T cells. **A)** UCSC browser tracks displaying ChIP-seq signals at the *Hhex* gene for endogenous PU.1 (αPU.1) in DN1-DN2b stages, and PU1WTHA and PU1ENGHA introduced into CD25^+^ cells cultured from fetal liver precursors (magenta tracks). Also shown are ATAC signals in the indicated stages (black tracks), H3K4me2 signals (red tracks), endogenous RNA peaks (purple tracks), and RNA in exogenous PU.1 transductants (green tracks). Highlight: region closing from the ThyDN1 to ThyDN3 stages with loss of endogenous PU.1 binding during normal transcriptional silencing from DN1 to DN3. **B)** Heatmaps show hierarchically clustered expression of genes affected by PU1WTHA transduction of DN2b/DN3 cells. Selected gene names are shown. Red side bars: genes with associated exogenous PU.1WTHA binding; black bars: genes with no associated peaks (proximity based; HOMER). Left block: DEGs defined in cells remaining CD25^+^ (PU1WTHA25). Right block: DEGs defined in cells becoming CD25^−^ CD44^+^ (PU1WTHA44)(right). Responses of these DEGs to constructs indicated below panels are compared. **C)** Comparison of site occupancy by endogenous PU.1 (DN1-DN2b, detected by αPU.1) and by exogenous PU.1 in parallel (PU1WTHA, detected by αHA) as a function of PWM score, at “open” and “closed” sites. Orange, open ThyDN1; blue, closed ThyDN1; green, open ThyDN3; gray, closed ThyDN3. **D)** Heatmap of binding of exogenous PU1WTHA, PU1ENGHA, and PU1ETSHA in CD25^+^ DN2b/DN3 primary cells, compared with endogenous PU.1 in DN1, DN2a, and DN2b, ATAC-seq in ThyDN1 and ThyDN3, and H3K4me2 -and H3K27me3 in DN1 and DN3 cells. Color scales: tag count densities. **E)** DN1 αPU.1 (top left), exogenous PU1WTHA αHA (bottom left) and ATAC (top right) distribution plots within 1 kb of endogenous or exogenous PU.1 bound sites from Groups 5, 6, and 7 in Fig. 5D. **F)** Association of Groups 5, 6, and 7 sites with changes in linked gene expression induced by exogenous PU1WTHA in CD25^+^ cells. *** K-S p-value ≤ 0.0001, ** p-value ≤ 0.001,* p-value ≤ 0.05.

First, the effects on genes linked to exogenous PU.1 occupancies were generally biased toward positive regulation: genes close to sites bound by PU1WTHA showed a trend of being slightly upregulated by PU.1 compared to genes without sites (Fig. S4E, Kolmogorov-Smirnov p ∼ 10^−6^). Concordantly, local binding by PU1ENGHA was linked with repression (K-S p ≤ 0.0001); the effects of PU1ENGHA were in good agreement with the shorter-term effects we reported previously (Champhekar et al. 2015)(Fig. S4F). The bias toward activation by linked PU1WTHA occupancy was robust among the 1586 differentially expressed genes responding significantly to PU1WTHA in CD25^+^ samples (DEGs, padj<0.1). Of these, 1368 were linked to new occupancy sites, and of these linked genes 1046 responded positively (Table S3). Fig. 5B shows heat maps of these patterns of expression of responding genes as defined in PU1WTHA25 (left panel, 1586 DEGs overall) or defined in PU1WTHA44 cells (right panel, 7965 DEGs overall), and separated into genes with (red line, right margin) or without (black line, right margin) linked sites of PU1WTHA occupancy. About 90% of DEGs in PU1WTHA25 cells were linked to sites of exogenous PU.1 binding (red margin), and most of these were upregulated (PU1WTHA25 vs. EV, column 2 vs. 1). Expression of the same genes was further increased in transductants becoming CD44^+^ CD25^−^ (Fig. 5B, left, column 5). As expected, the obligate repressor PU1ENGHA (column 3) downregulated many of the PU1WTHA-induced genes with binding sites, and had little effect on those without direct PU.1 binding; the PU1ETSHA construct (column 4) had limited impact. These results are consistent with PU.1 acting primarily as an activator of genes responsive to its direct binding.

### PU.1 activates pro-T cell target genes via developmentally dynamic non-promoter sites

Exogenous PU1WTHA added to DN2b/DN3 cells quantitatively restored occupancy of genomic sites to levels similar to those in normal DN1 cells, with an almost indistinguishable distribution of site quality preferences as compared to earlier endogenous PU.1 based on PWM scores (Fig. 5C). Globally as at *Hhex* (Fig. 5A), the specific sites occupied by exogenous PU1WTHA overlapped those occupied by endogenous PU.1 in earlier stages (Fig. 5D) and exhibited similar properties in terms of motif enrichments (Fig. S5A; cf. Fig. 1G). Sites of exogenous PU.1 binding with ATAC and histone modification properties similar to endogenous PU.1 sites of both Groups 1 and 2 (cf. Fig. 2A) were engaged. However, PU1WTHA in transduced CD25^+^ cells also entered some sites that had not been occupied by endogenous PU.1 previously. Three classes of sites could be defined (Fig. 5D, Groups 5, 6, 7), based on their histories of normal endogenous PU.1 binding at earlier stages, ATAC accessibilities, and histone marks in normal DN1-DN3 cells. The Group 5 sites were highly overlapping with Group 2 sites bound by endogenous PU.1 in DN1 and DN2a stages. They had reduced ATAC accessibility and H3K4me2 marks between DN1 and DN3 stages (Fig. 5D,E), like the highlighted sites in *Hhex* and *Itgam* (Fig. 5A; Fig. S5B, lower). Group 7 sites were constitutively open (Fig. 5D,E), exemplified by sites in *Il2ra* (Fig. S5B, upper). These tended to have lower PWM scores than Group 5 sites (Fig. S5C) and unusually high co-enrichment of CG-rich motifs (Fig. S5D). Again, these Group 7 sites corresponded to the promoter-enriched sites of endogenous PU.1 binding (cf. Fig. 2A, group 1). The new category of sites bound by exogenous PU1WTHA (Fig. 5D, Group 6) showed no obvious history of significant binding by endogenous PU.1 earlier, and consisted of sites that were normally ATAC-”closed” and poorly marked with H3K4me2, at DN1 and DN3 stages alike (Fig. 5D,E). Group 5 and Group 6 sites both had high quality PU.1 sites, but with slightly higher PU.1 site scores for Group 6 sites and substantially higher enrichment of Runx motifs in the physiologically occupied Group 5 sites (Fig. S5C,D). The ability to occupy these closed, “ectopic” Group 6 sites thus most likely resulted from the slightly elevated concentration of the exogenous PU1WTHA in these transduced CD25^+^ cells. However, at both Group 5 and Group 6 sites, the exogenous PU.1 showed its ability to enter chromatin sites that were closing or frankly closed in these pro-T cells.

The impacts of exogenous PU.1 binding on transcription were strikingly different among genes linked to these different classes of sites, as judged by stratifying DEGs based on site classes (Fig. 5F). The contributions of different site types were additive when linked to the same genes (Fig. S5E). Among DEGs with sites falling into only one site class, exogenous PU1WTHA preponderantly upregulated genes linked only to Group 5 sites (Fig. 5F, blue). It also upregulated those with Group 6 sites only (Fig. 5F, green), despite their normally closed status. In contrast, genes with PU1WTHA binding only at Group 7 sites showed slight or negative effects on average (Fig. 5F, red). The DEGs with these constitutively open, promoter-enriched sites increased expression in PU1WTHA25 cells even less than “background” genes with no PU1 binding (Fig. 5F, black; p<0.05), or were even slightly downregulated, especially in cells that became CD44^+^ (Fig. S5E lower panel, red vs. black, p<0.0001). These results confirm that PU.1 binding most commonly works to activate genes in pro-T cells, and does this primarily through non-promoter sites where the chromatin is not constitutively open.

### Signatures of PU.1-repressed gene sites

Although less frequent than positively regulated genes, the highest-confidence PU.1 target genes also included some genes repressed by PU.1 that also had local binding sites (Fig. 4H, Table S6A). To obtain better enrichment for candidate repressive sites than was possible in Fig. 5D, we compared two groups of PU1WTHA occupancy sites, distinguished by dynamic ATAC accessibility changes in normal development, and separated into those that increased and those that lost ATAC accessibility in the transition from DN1 to the DN3 stage, when endogenous PU.1 decreases (Fig. 6A). As this analysis excluded constitutively open sites, >95% of both opening and closing sites were distal. Binding sites linked with naturally PU.1-activated genes were expected to become less accessible from DN1 to DN3 (closing), as for Group 5 sites (Fig. 5D,E) that are mostly positively regulated by PU.1WTHA. Correspondingly, if any genes were directly repressed by endogenous PU.1, their PU.1 binding sites could be opening during this transition. Both opening and closing sites were similarly co-enriched for PU.1 and Runx motifs, although endogenous PU.1 binding was more common at the closing sites (5077/6888) than at the opening sites (1792/4283). However, the closing sites showed some enrichment for composite PU.1-IRF motifs, while the opening sites were enriched for E protein (Tcf3) and Tcf7 family motifs (Fig. 6B).

**Figure 6:**
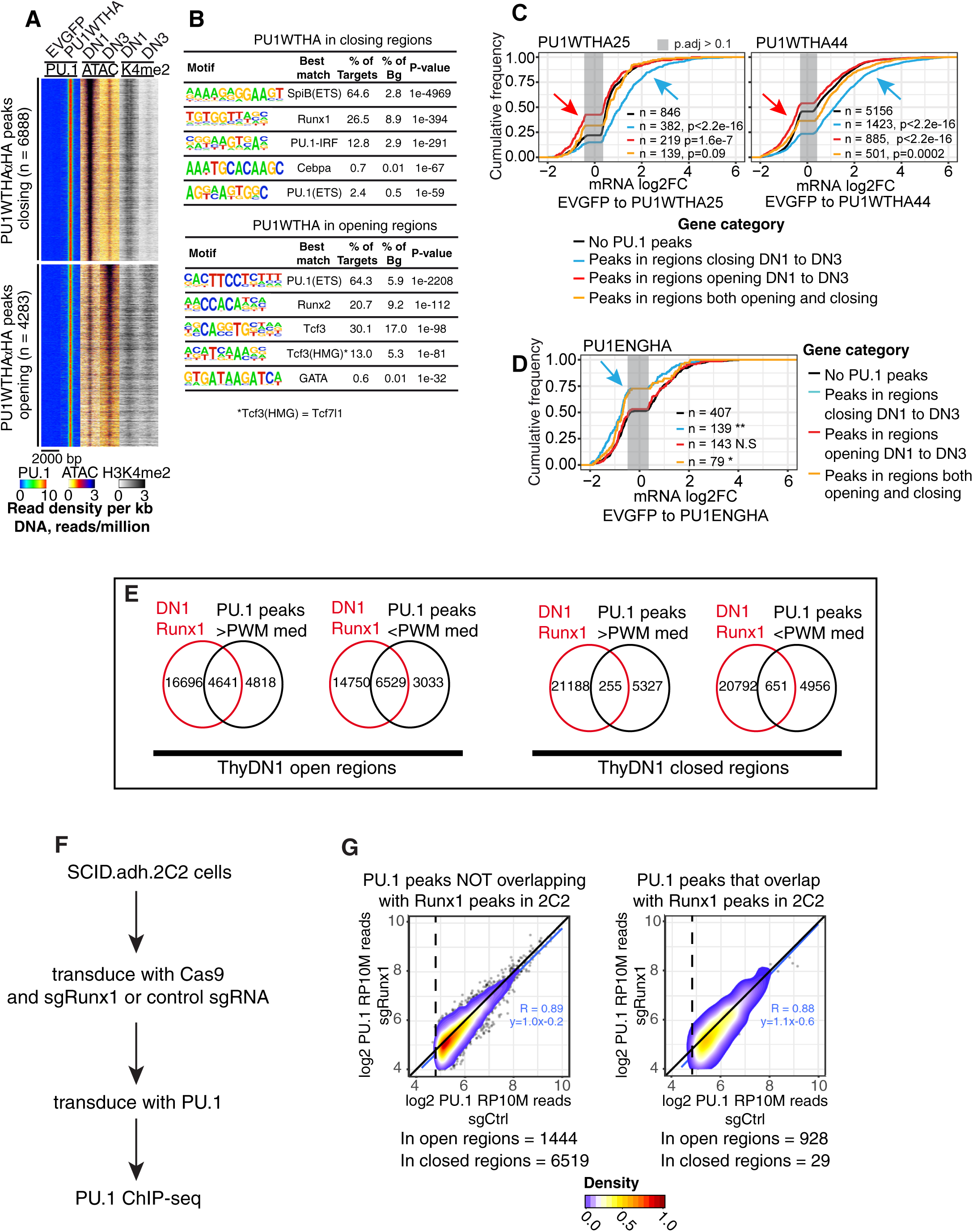
Dynamic changes in ATAC accessibility in primary pro-T cells predict functionality of PU.1 binding. **A)** Definition of sites of exogenous PU1WTHA binding in transduced CD25^+^ cells at sites that are naturally closing (top) or opening (bottom) *in vivo* from DN1 to DN3 stages. Heat maps show exogenous PU1WTHA (αHA) ChIP-seq, ATAC-seq, and H3K4me2 tag count distributions. Peaks are ordered by their PU1WTHA tag counts (high to low). **B)** Top five motifs enriched at PU1WTHA αHA peaks in regions that change dynamically from ThyDN1 to ThyDN3. **C)** Changes in gene expression induced by acute PU1WTHA introduction in genes linked to sites that would normally close (blue) or open (red) from ThyDN1 to ThyDN3, as compared to genes with no PU1WTHA peaks or genes with a mixture of opening and closing peaks. Left: genes with significant differential expression induced in cells remaining CD25^+^. Right: genes with significant differential expression induced in cells becoming CD44^+^. *** K-S p-value ≤ 0.0001. **D)** Effects of PU1ENGHA on genes linked to sites in opening or closing regions. Arrow shows that obligate repressor effect is most strongly correlated with sites that are only open when endogenous PU.1 is expressed. **, K-S p-value ≤0.001. **E)** Venn diagrams of overlaps between Runx1 binding in primary DN1 cells (Hosokawa et al. 2018) and PU.1 occupancy sites in primary DN1 cells, stratified by their ATAC status and PWM scores above or below median. **F)** Schematic of experimental test for Runx1-dependence of exogenous PU.1 site binding in Scid.adh.2C2 cells. PU.1 in the pMXs vector was used. **G)** Results of protocol shown in **F**. PU.1 occupancy scores for PU.1 binding in cells after Runx1 disruption (y axis) are plotted against scores for PU.1 binding in controls (detected with αPU.1). Sites are stratified according to their ATAC accessibility and Runx1 occupancy in unmanipulated Scid.adh.2C2 cells (Hosokawa et al. 2018). Pearson’s r and linear trend lines are shown for peaks with a peak score (Homer *findPeaks*) >30 in controls.

The highest-confidence naturally PU.1-regulated target genes (Table S6; Fig. 4H) were indeed enriched for the presence of such linked dynamic PU.1 binding sites (Fig. S6A,B). PU.1-activated genes were enriched near sites that closed in development (p<10^−6^), while the rare PU.1-repressed genes were enriched near sites that opened (p<10^−4^). Across the genome, these criteria identified sites linked to genes that respond robustly to PU1WTHA binding, and enabled repression targets to be substantially enriched (Fig. 6C). PU1WTHA binding at non-promoter sites that normally close from DN1 to DN3 stage was associated with “re-activation” of linked genes, as compared to genes with no or mixed peaks (Fig. 6C, p<2.2×10^−16^). It had little effect on repression (blue curve shifted from others more on the right than on left). In contrast, binding to regions that open up naturally in cells progressing from DN1 to DN3 (Fig. 6C, red curves) yielded enhanced repression relative to genes lacking PU.1 binding sites, without increased activation (red curve shifted on the left, not on the right). Genes linked to developmentally closing sites were more likely both to be activated by full-length WT PU.1 and to be repressed by obligate repressor PU1ENGHA (Fig. 6D). Thus, dynamic distal sites, marked by distinct co-enrichments of partner motifs, were preferential targets for endogenous and exogenous PU.1 to mediate positive or negative regulation.

### PU.1 can bind its target sites independently of Runx1

The motifs most highly co-enriched with PU.1 motifs at both activating and repressing sites were for Runx factors. In a related study, we have shown that PU.1 physically interacts with Runx1 in pro-T cells and recruits it to PU.1 binding sites (Hosokawa et al. 2018). Fig. 6E shows that a large fraction of endogenous PU.1 occupancy sites that were open in primary DN1 cells were in fact co-occupied by Runx1 [data from (Hosokawa et al. 2018)], with highest Runx1 binding enrichment among those sites with lower-quality PU.1 motifs (below median PWM score). Runx1 co-occupancy was much lower at PU.1 sites in closed regions (Fig. 6E) consistent with motif enrichments (Fig. 1G, Fig. S1D). These results raised the question of how much Runx1 was contributing to the ability of PU.1 to establish its own binding at different classes of sites.

Whereas primary cells are severely inhibited by combined deletion of Runx1 and forced expression of PU.1, we could test the effect of Runx1 loss on PU.1 binding using exogenous PU.1 in the Scid.adh.2C2 cell system, as shown in Fig. 6F (see Methods). Fig. 6G compares PU.1 ChIP-seq signals at non-promoter sites in Scid.adh.2C2 cells that had also previously been transduced with Cas9 plus either control sgRNA (sgCtrl) or sgRNA against Runx1 (sgRunx1). This treatment effectively disrupted the region around the first Met codon shared by all Runx1 isoforms (Fig. S6C) and eliminated Runx1 protein expression (Hosokawa et al. 2018). As for endogenous PU.1 in primary cells, exogenous PU.1 occupancies overlapped pre-existing Runx1 sites (right panel)(Hosokawa et al. 2018) much more in regions that were initially open in 2C2 cells than in regions that were closed. However, high-quality PU.1 binding to non-promoter sites was essentially unchanged by Runx1 disruption (diagonals in Fig. 6G), whether the sites overlapped pre-existing Runx1 peaks (right) or not (left). Thus, although Runx1 can be important for PU.1 function in pro-T cells (Hosokawa et al. 2018), PU.1 binding *per se* is not Runx1-dependent.

### PU.1 requires its non-DNA binding domains to occupy sites in closed chromatin

PU.1 in pro-T cells fulfills three criteria for pioneering, as it uses its own DNA binding affinity to select binding sites in closed and open chromatin, it maintains accessibility of open sites and can open chromatin at many closed sites, and its most common binding partner, Runx1, is not required for PU.1 binding in either type of site. However, the question remained whether PU.1’s own DNA binding specificity was sufficient to determine its sites of action. To address this, we mapped the binding sites for the isolated PU.1 DNA binding domain alone (PU1ETSHA) (constructs diagrammed in Fig. S7A). Both PU1ETSHA and its derivative, PU1ENGHA, could bind to DNA in primary CD25+ pro-T cells and in Scid.adh.2C2 cells with site quality criteria similar to those of PU1WTHA (Fig. S7B,C). However, when PU1WTHA, PU1ETSHA, and PU1ENGHA were introduced into Scid.adh.2C2 cells, despite strong expression of PU1ETSHA especially (Fig. S7D), there were dramatic, site-specific differences between the binding of both truncated constructs and that of full length PU.1 (Fig. S7E). At open sites, with strong ATAC and H3K4me2 signals in non-transduced Scid.adh.2C2 cells, the two truncated constructs bound similarly to full-length PU1WTHA. These were often sites occupied by another ETS factor, Ets1 (Fig. S7E), in parental Scid.adh.2C2 cells. But at sites that were initially closed in non-transduced Scid.adh.2C2, PU1ENGHA and PU1ETSHA showed much less binding than PU1WTHA. Successful binding by PU1ETSHA and PU1ENGHA was accordingly biased toward promoters, especially in Scid.adh.2C2 cells (Fig. S7F).

The same sites that were relatively poorly bound by PU1ETSHA and PU1ENGHA in the cell line were not poor quality based on sequence, for in the primary pro-T cells these sites (Fig. S8A,B, Group A sites) were bound by these constructs as well as the sites that were accessible in the Scid.adh.2C2 cells (Group B sites). Group A sites were sites of normal endogenous PU.1 binding (Fig. S8A,B), and in primary cells, where both site groups were accessible, the Group A sites were far better at mediating the repressive effect of PU1ENGHA (Fig. S8C) than the constitutively open, promoter-enriched sites (Group B). Fig. 7A-D show profiles of genes with such differential binding (highlighted) at Group A type sites. *Cd34, Syk, Robo1* and *Ffar2* are positively regulated by full-length PU.1 and repressed by PU1ENGHA in primary cells (Table S3). Additional genomic loci with similar occupancy patterns, *Cd44, Vav1, Elovl5, Flt3, Notch2*, and *Myd88*, are shown in Fig. S9.

**Figure 7:**
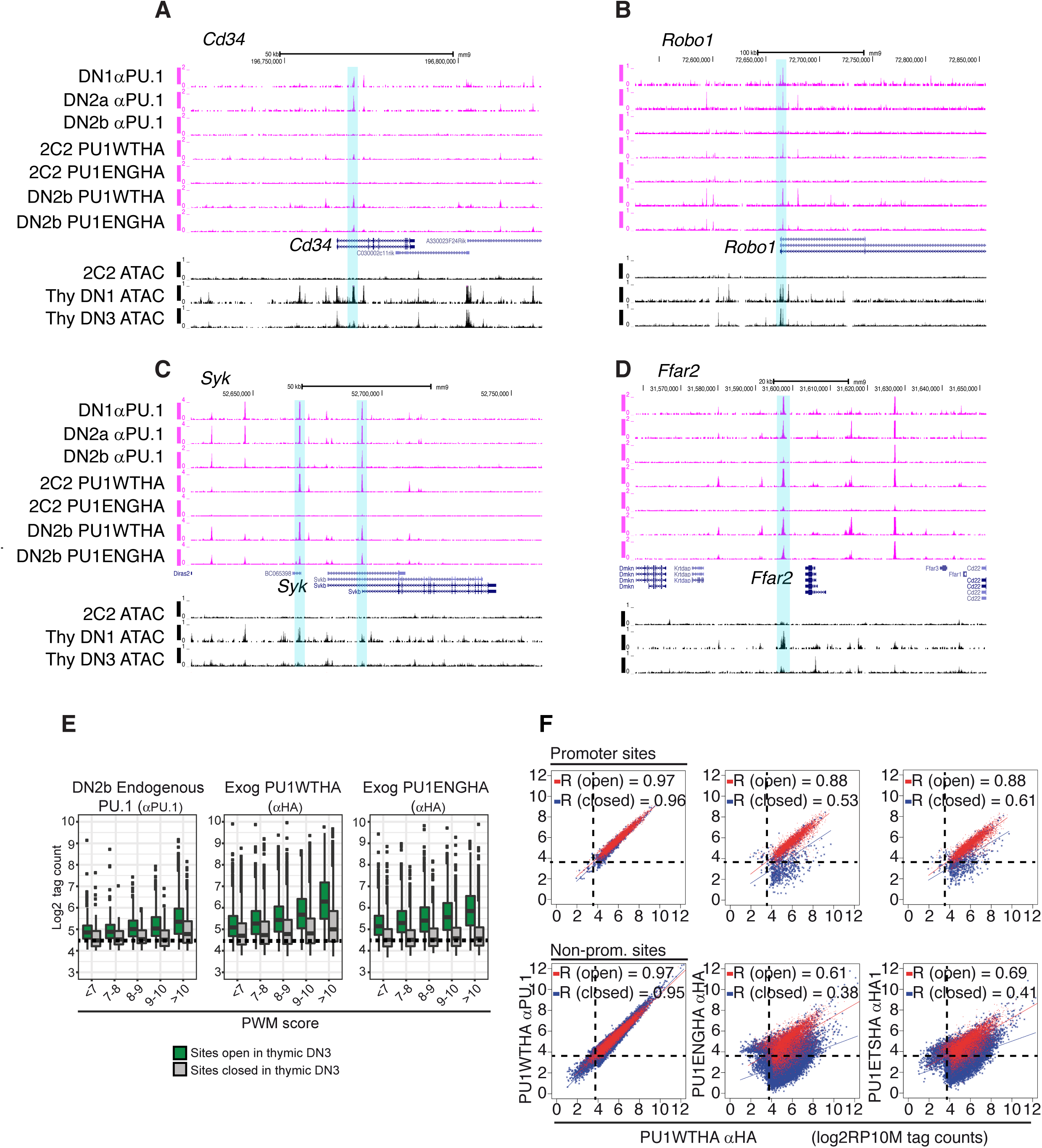
Full-length PU.1 is required for access to closed chromatin. **A)-D)** UCSC browser tracks comparing binding of endogenous PU.1, full-length exogenous PU1WTHA, and PU1ENGHA to open and closed regions in primary DN2b/3 CD25^+^ cells and Scid.adh.2C2 cells. Shown are the genes: **A)** *Cd34*, **B)** *Robo1*, **C)** *Syk***, D** *Ffar2***. E)** Comparison of PU1ENGHA binding with that of PU1WTHA and baseline binding of endogenous PU.1 in primary DN2b cells. Analysis as in Fig. 5C. Green bars: open sites in primary DN2b-DN3 cells. Gray bars: closed sites. Dashed line: background. **F)** Comparison of exogenous PU1WTHA with PU1ENGHA or PU1ETSHA binding in transduced Scid.adh.2C2 cells at open and closed promoter and non-promoter sites. PU1WTHA αHA counts in open (red) and closed (blue) regions are plotted against PU1WTHA αPU.1 (left), PU1ENGHA αHA (middle) or PU1ETSHA αHA (right) tag counts in promoters (top three panels) or non-promoter genomic sites (bottom three panels). n, peaks at open promoters=4498; n, peaks at closed promoters=810; n, peaks at open non-promoter elements=8719; n peaks at closed non-promoter elements=33158. Pearson’s r shown. Dashed lines: tag threshold for peaks considered bound.

The failure of the isolated PU.1 DNA binding domain (or its Engrailed derivative) to interact with these sites in the context of the DN3-like cell line suggested that PU.1 without its native trans-activation and protein-interaction domains could be more sensitive to chromatin state than full-length PU.1 for access to its binding sites. We tested whether a qualitative requirement for open chromatin could be distinguished from alternatives, either that the truncated constructs have lower affinity or that they reach a lower effective protein concentration. First, we compared binding success of PU1WTHA and PU1ENGHA in primary CD25+ cells at sites of different PWM quality scores with open vs. closed ATAC status. Here, PU1ENGHA bound comparably to PU1WTHA at open sites (Fig. 7E). However, whereas PU1WTHA showed some affinity cost for entry into closed chromatin, the data suggested much less binding by PU1ENGHA at closed sites, even when they contained very high-quality motifs (Fig. 7E). Wildtype and mutant constructs also showed sharply different binding titrations against target sites in SCID.adh.2C2 cells depending on whether they were ATAC-open or ATAC-closed. Fig. 7F shows the occupancies of the same open and closed sites by PU1WTHA, PU1ENGHA, or PU1ETSHA, in each case compared to PU1WTHA (detected by a different antibody). In these graphs, identical binding falls on the diagonal, as shown when the same PU1WTHA was detected with αPU.1 or with αHA antibodies. If there were global DNA-binding efficiency differences between a truncated construct and the full-length form we would expect to see the whole trend line shifting downward on the log/log plot or else changing slope. Instead, Fig. 7F shows a split pattern for the truncated constructs. The binding patterns of both truncated constructs closely matched PU1WTHA on open sites, but were specifically penalized, within the same cells, with lower binding slopes and high dispersions at closed sites. Thus, the non-DNA binding domains of PU.1 strongly and selectively affected the ability of the PU.1 DNA binding domain to establish occupancy in closed chromatin.

These results demonstrate that non-DNA binding domains of PU.1 not only affect the functional output of the binding, but also affect the ability of the factor to engage with novel, unoccupied distal sites where accessibility has not been established. The full site-selectivity of PU.1 as a pioneer, therefore, depends not only on its ability to find high-affinity sites, but also on the ability of its protein-interaction domains to convert recognition of sites in closed chromatin into stable, functional occupancy.

## DISCUSSION

PU.1 is known to act as a pioneer in the role of a lineage determining factor that initiates and sustains identity of myeloid cells. One question in this study has been whether its inherited high activity in the earliest pro-T cells also shapes the epigenetic landscape in this context, even though its activity in this lineage is hit and run and many of its known partners are not expressed. Criteria for pioneering involve the ability to open closed chromatin and to initiate occupancy of a cis-regulatory site by a complex of transcription factors (Zaret and Carroll 2011). Logically, this implies that pioneer factors can (1) find and bind to sites in closed chromatin; (2) bind to target sequences based on their own specificity, even before other factors are available to collaborate; and (3) initiate the opening of chromatin as new partners are recruited. In the myeloid system, PU.1 can play these roles (Natoli et al. 2011; Barozzi et al. 2014) but often it does so as part of a partnership with C/EBP family factors, where either PU.1 or the C/EBP factor can help nucleate binding by the other (Laiosa et al. 2006; Feng et al. 2008; Heinz et al. 2010; Heinz et al. 2013). As we show here, PU.1 does fulfill most of the criteria for pioneering in pro-T cells. It can bind closed chromatin as well as open. Its binding is primarily guided by its own well-defined specificity, even though we show that binding stably to closed chromatin sites in these cells requires a consistently higher affinity of site recognition than binding to open chromatin. Its binding preferences follow its own concentration, with its binding persisting only on higher and higher affinity sites as its concentration decreases, consistent with mass-action in which binding is dominated by PU.1’s own specificity. Finally, it can clearly trigger chromatin opening, and it is required continuously to maintain accessibility at many of its target sites. These functions are exerted especially at a major subset of its non-promoter sites, where PU.1 can rapidly induce transposase accessibility followed by recruitment of histone acetyltransferases. It may cooperate with Runx1 in causing chromatin to open and to activate its positive target genes (Hosokawa et al. 2018), but it does not require Runx1 for its own binding. These findings are summarized in Fig. 8.

**Figure 8:**
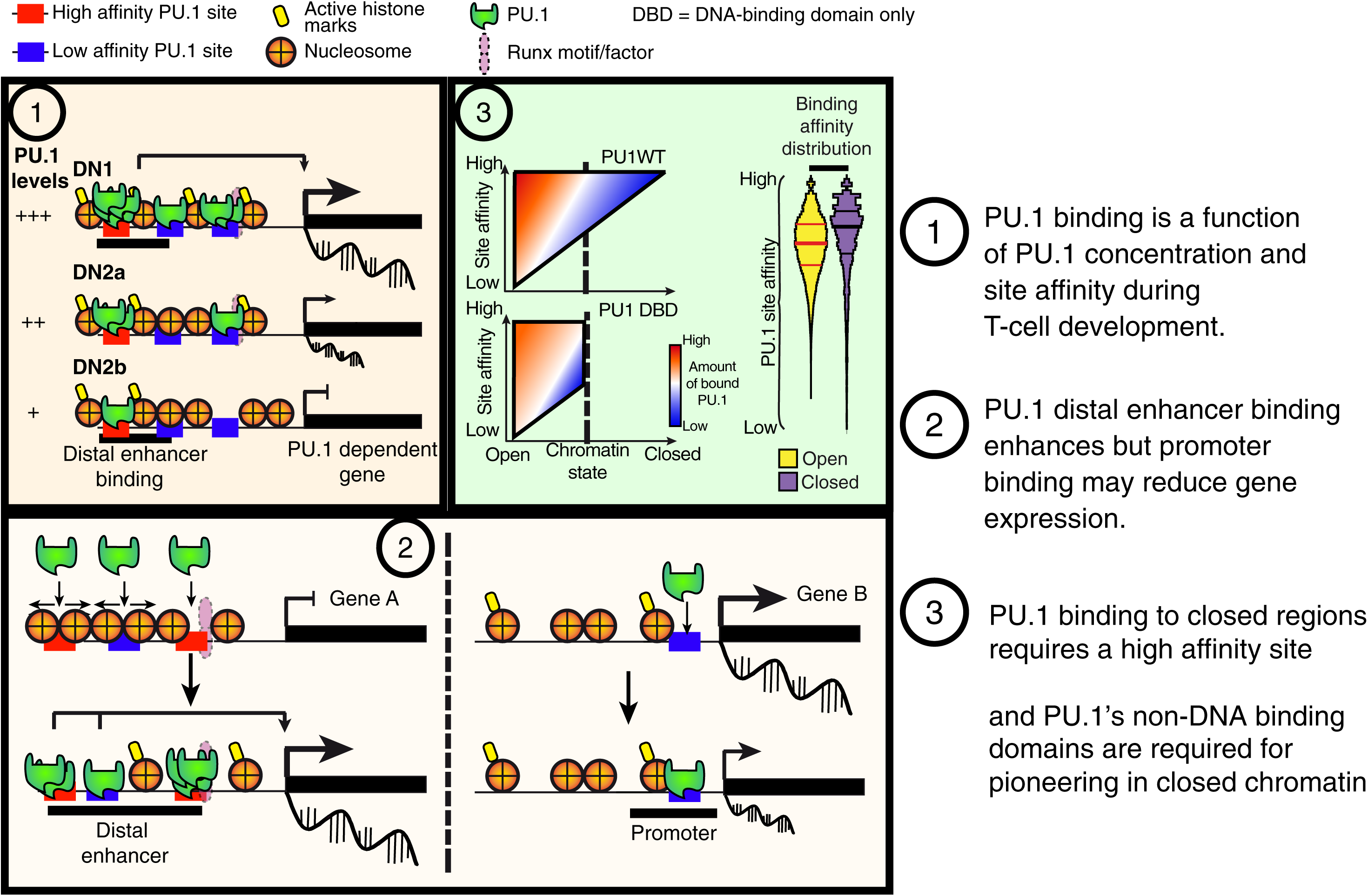
Schematic of PU.1 site choice criteria and functional impacts in pro-T cells. Summary of mechanisms regulating PU.1 binding and function (see Discussion).

By bringing together ChIP-seq binding analysis, short-term forced expression and acute, synchronized Cas9-mediated deletion tests, the positive regulatory targets of direct PU.1 binding in early pro-T cells have now been defined robustly. A core group of about 250 genes is downregulated by PU.1 disruption in pre-commitment pro-T cells and upregulated by addition of PU.1 to post-commitment pro-T cells, and these genes are linked to non-promoter sites of PU.1 binding that depend on PU.1 for ATAC accessibility and lose accessibility when PU.1 levels decline. On the other hand, a minority of non-promoter regulatory elements that bind PU.1 increase accessibility during commitment, and these are enriched for sites where PU.1 binding directly mediates repression. These high-confidence target genes of PU.1, both positive and negative, have direct binding, chromatin features at PU.1 sites and developmental expression implying that normal endogenous PU.1 activity is a major contributor to their developmental profiles within the context of the pro-T cell program itself.

The high developmental resolution possible in this system sheds light on features of PU.1 action that have much broader implications. The first is that motif quality of sites, magnitudes of occupancy, promoter-association, and immediate degrees of ATAC accessibility are useful but not sufficient criteria to identify loci of actual PU.1 regulatory action. Promoters are by far the most accessible sites for PU.1 binding: not only are they constitutively open as measured by ATAC-seq, but also they offer an exceptionally permissive affinity threshold for PU.1 engagement, and they do not require the PU.1 transactivation domains for efficient PU.1 binding. In myeloid cells, PU.1 is commonly bound at promoter-proximal regulatory elements of functional target genes (Gonzalez et al. 2015). However, although PU.1 is frequently bound to promoters of active genes in pro-T cells as well, in this cell context the actual functional impact of its activity at these promoters is often weak or null, unless supplemented by local binding to non-promoter sites as well. Instead, it is those dynamic distal elements, even those with less-pronounced PU.1 binding, that are best associated with function.

PU.1 binding is excluded by H3K27me3-marked repressive chromatin (Zhang et al. 2012), but modestly elevated PU.1 can also enter other regions that appear “inaccessible” by criteria of ATAC-seq, and upon binding can mediate rapid chromatin opening and local gene activation. We have shown that the ability to establish binding in such closed sites depends on a function of PU.1 structurally distinct from its site recognition. Although the PU.1 DNA binding domain alone binds to the genome with a motif preference matching that of full-length PU.1, its pioneering potential is much constrained due to a qualitative exclusion from closed sites. Like the case of EBF1 (Boller et al. 2016), whereas the PU.1 DNA-binding domains define its nucleotide sequence specificity, the non-DNA binding domains of PU.1 are required for its access to closed sites. Thus, the DNA binding domain alone is not only deprived of discrete transactivation or protein interaction functions, but also of access to a large subset of potential genomic targets. This effect explains some of the functional specificity of the obligate-repressor PU1ENGHA form as previously reported (Champhekar et al. 2015), which can repress competitively at endogenous PU.1 binding sites as long as they are open but cannot gain access to those sites once they have developmentally closed. The non-DNA binding domains of PU.1 are important for its interactions with protein partners, at least including Runx1 and Satb1 in early pro-T cells (Hosokawa et al. 2018), as well as factors important in erythroid development (Zhang et al. 2000; Stopka et al. 2005). Although Runx1 itself may not explain this specific binding requirement, many other interacting proteins (Hosokawa et al. 2018) remain as candidate collaborators to assist PU.1 binding in closed chromatin of pro-T cells.

Thus, despite PU.1’s fulfillment of simple criteria of pioneer function, its impact on early T-lineage cells is determined by the intersection of its own powerful binding activity and its dynamic interaction with other determinants of epigenetic state. The effect of closed chromatin on full-length PU.1 appears to be quantitatively compensated by the higher binding affinities at nearer-optimal sites, but the transcriptional effect of this PU.1 binding on neighboring genes depends on other factors that determine whether the site bound is constitutively open, constitutively closed, or may be dynamically opening or closing in development. Furthermore, the restrictions on the PU.1 binding domain itself imply that PU.1’s pioneering activity also depends on interactions with other transcription factors or chromatin modulators. This reconciles the biochemical pioneering-like activities of PU.1 with the distinctive lineage-specific occupancy pattern of its activity in the earliest stages of T-cell development.

## METHODS

Cell purification, cell lines, culture conditions, retroviral constructs and transduction, Cas9 deletion, RNA-seq and ChIP-seq methods were similar to those in (Hosokawa et al. 2018), adapted from previous work (Zhang et al. 2012; Del Real and Rothenberg 2013; Champhekar et al. 2015). All sample types were generated in 2-4 separate biological replicates, with inter-sample correlations as shown in Fig. 4 and Fig. S10. Detailed methods including statistics and bioinformatics are presented in Supplemental Information.

## Data access

Accession numbers for data deposited in GEO:GSE93755 and GSE110020.

## Acknowledgments

We gratefully acknowledge advice on specific algorithms from Jeffrey Longmate (Beckman Research Institute of the City of Hope) and Hao Yuan Kueh (Caltech and University of Washington), and generous advice and discussion from Barbara Wold (Caltech). We thank Marissa Del Real, Ameya Champhekar, and members of the Rothenberg group for constructs and helpful discussion, Diana Perez, Jaime Tijerina, and Rochelle Diamond for cell sorting and advice, Ingrid Soto for mouse colony care, Vijaya Kumar for library preparation and sequencing, Henry Amrhein and Diane Trout for next-generation computational assistance and Igor Antoshechkin for sequencing facility management. This research was supported by fellowships from the Swedish Research Council (to JU), from the Manpei Suzuki Diabetes Foundation (to HH), grants from the USPHS to EVR (R01HD076915 and R01AI95943), by the L. A. Garfinkle Memorial Laboratory Fund and the Al Sherman Foundation, and by the Albert Billings Ruddock Professorship to EVR.

